# NeotropMyrtaceaeFruits: a database of fleshy fruits and seeds of Neotropical Myrtaceae

**DOI:** 10.64898/2026.11.24.768853

**Authors:** Hercília Freitas da Cunha, Yacov Kilsztajn, Jair Eustáquio Quintino de Faria Júnior, Thiago Fernandes, Paulo Henrique Gaem, Marla Ibrahim Uehbe de Oliveira, Carolyn Elinore Barnes Proença, Aline Stadnik, Karinne Sampaio Valdemarin, Fábio Christiano Speck Vieira, Vanessa Graziele Staggemeier

## Abstract

Myrtaceae is one of the most species-rich plant families in the Neotropics, but we still don’t know which factors explain its high success in this region. Databases containing information on species traits are essential to understand lineage diversification and their evolutionary success. Some studies indicate that flower morphology is highly homogeneous within the family in the Neotropics and offers limited insight into species diversification. Alternatively, fruit and seed morphology remain poorly explored and could provide valuable information. The wide morphological variation in fruits and seeds within this family is remarkable, ranging from species with fruits smaller than 0.5 cm to those exceeding 10 cm, and from fruits containing a single small or large seed to those with hundreds of tiny seeds. This diversity has probably played a key role in the group’s evolutionary history, particularly through interactions with seed dispersers. However, information on fruit morphology is scattered throughout the literature, including articles with taxonomic descriptions, floras and ecological studies, as well as unpublished literature such as dissertations and theses. To fill this gap, we combined information from the scientific literature with the expertise of Myrtaceae specialists to compile a database on fruit and seed length and diameter, seed number, fruit color, and display type (unicolored vs. multicolored) for Neotropical species. We included taxonomic information in the database (such as subtribal classifications and infrageneric sections for the large clades *Eugenia, Myrcia*, Pliniinae and *Psidium)* to enhance its potential use in macroecological studies. Through active searching, 250 references were consulted, of which 216 were scientific articles, 15 books, 17 dissertations and theses and 2 virtual databases. We also collected new data from the field and herbarium to complement the dataset. The resulting database includes information from 1997 records for 997 species. This database provides valuable data for analyzing evolutionary trends, helping to explain the patterns of clade diversification. It also offers a foundation for future studies on the ecological strategies of this highly diverse family, such as understanding the ecological interactions with the disperser fauna and the role of abiotic factors shaping the diversity of fruit and seeds in the Neotropics. This database is licensed under the Creative Commons Attribution 4.0 International License (CC BY 4.0). When using this database, please cite this datapaper.

## Overview

Myrtaceae is one of the most diverse plant families in the World, comprising approximately 130 genera and 6,292 species (POWO, 2026). Due to its high species richness and broad geographic distribution, the family has been considered a model group for investigating patterns of diversity and distribution in tropical floras (Lucas & Bünger 2015; Murray-Smith et al. 2009). Within the Neotropics, the family is represented mainly by the Neotropical clade of the tribe Myrteae (*sensu* Vasconcelos et al. 2017; hereafter Neotropical Myrtaceae), which comprises about 2,500 species and represents a large radiation of fleshy-fruited plants in the region (NMWG et al. 2024), providing an excellent system for comparative ecological and evolutionary studies. The objective of this study was to develop a comprehensive database of fruit and seed traits for Neotropical Myrtaceae species, which will allow future studies on macroecological and macroevolutionary perspectives. In short, the database compiles morphological information on fruits and seeds through an extensive review of the scientific literature, complemented by data obtained from herbarium specimens and field observations.

## Data Availability

The database will be made available on Figshare upon manuscript acceptance under the Creative Commons Attribution 4.0 International License (CC BY 4.0). Users of the database are requested to cite this data paper. The code used to calculate species-level mean values from the database will also be made available on Figshare.

## Data Description

The database compiles fruit and seed traits for Neotropical Myrtaceae species, combining information from the literature, field observations, and herbarium specimens, and reviewed by experts on the group. Variables include fruit and seed length and diameter, fruit color, number of seeds, fruit display type (unicolored or multicolored), plant height, geographic coordinates, data-collection method, and the original reference. Taxonomic information was subsequently added to assign each record to the appropriate subtribe and infrageneric section.

The database comprises 1,997 records gathered from 250 literature sources alongside new field and herbarium observations compiled in this study. Literature sources (*n* = 1,722 records) included 216 scientific articles (*n* = 1,025), 15 books (*n* = 321), 17 dissertations and theses (*n* = 260), and two online databases (Flora e Funga do Brasil 2026, *n* = 84; Jordano 2007, *n* = 5). Additionally, this study contributed 302 new records from primary field surveys and herbarium collections.

Collection methods for fruit and seed data were standardized as follows: “Field” (*n* = 156, 7.8%), for measurements taken from fresh material in the field; “Herbarium” (*n* = 1121, 56.1%), for measurements based on preserved specimens; “Field or herbarium” (*n* = 435, 21.8%), when both methods were cited but could not be distinguished; and “Untraceable origin” (*n* = 285, 14.3%), for data from secondary sources whose original measurement sources could not be determined. For Bufalo et al. (2016) and Bello et al. (2017), original sources were consulted whenever possible.

Mean species values were calculated using a custom R script, which is publicly available alongside the database. The NeotropMyrtaceaeFruits database comprises data for 997 species, encompassing 13 varieties of six species, and 29 genera, representing 40.4% of the approximately 2,500 species in the tribe Myrteae (Fig. 1). Specifically, the database contains data on fruit length for 742 species, fruit diameter for 934 species, seed length for 523 species, seed diameter for 446 species, and seed number for 661 species (Fig. 2). Data on fruit color and display are available for 748 and 404 species, respectively (Fig. 3).

**Figure 1.**
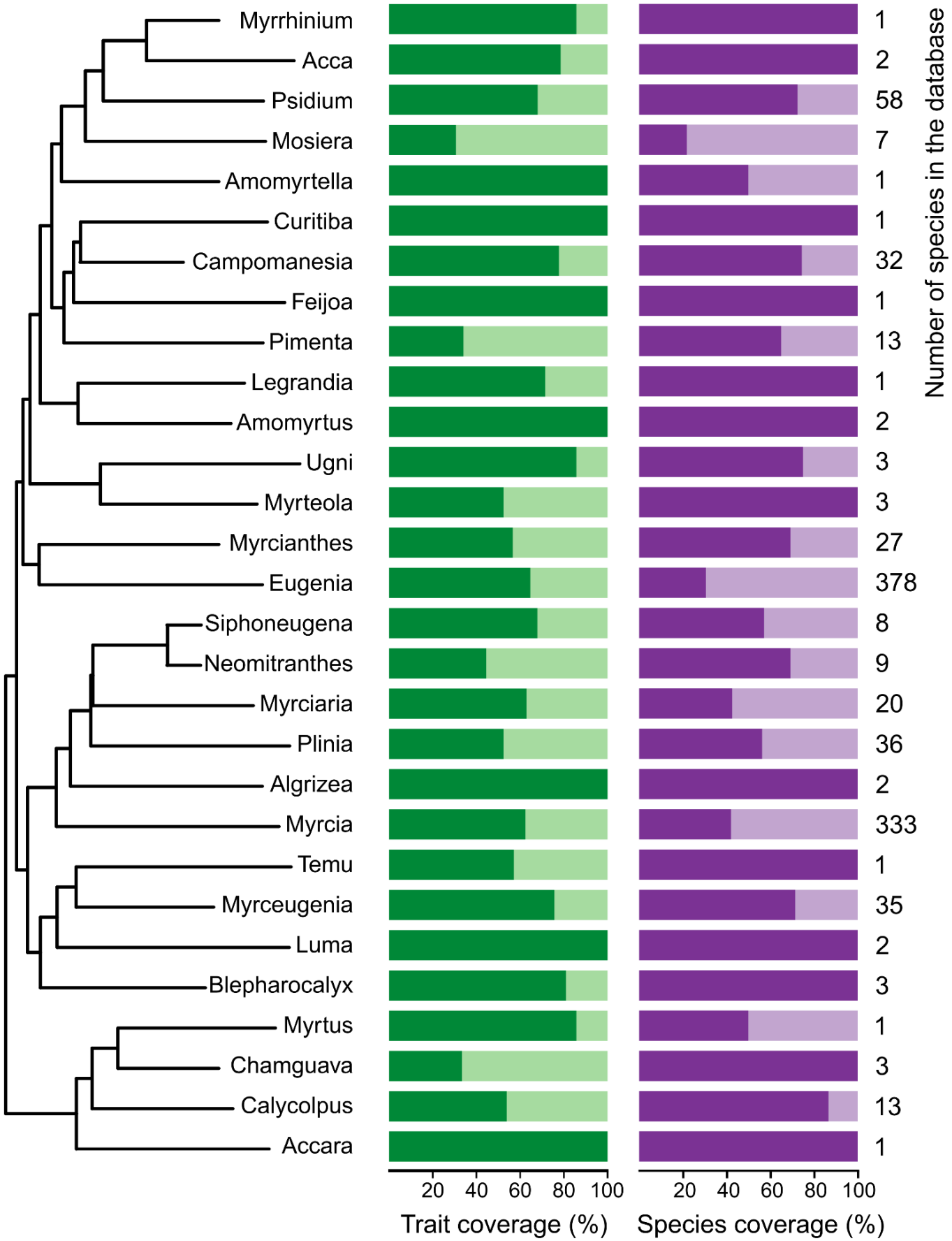
Trait and species coverage by genus. Trait coverage represents the percentage of possible species–trait combinations covered by the database, calculated as the number of available trait records divided by the total number of possible records (number of species in the database × seven traits: fruit diameter, fruit length, seed diameter, seed length, seed number, fruit color, and fruit display). Species coverage was based on WCVP for the Neotropical clade of the tribe Myrteae, and numbers on the right indicate the number of species included in the database. Genus-level phylogeny is adapted from NMWG et al. (2024).

**Figure 2.**
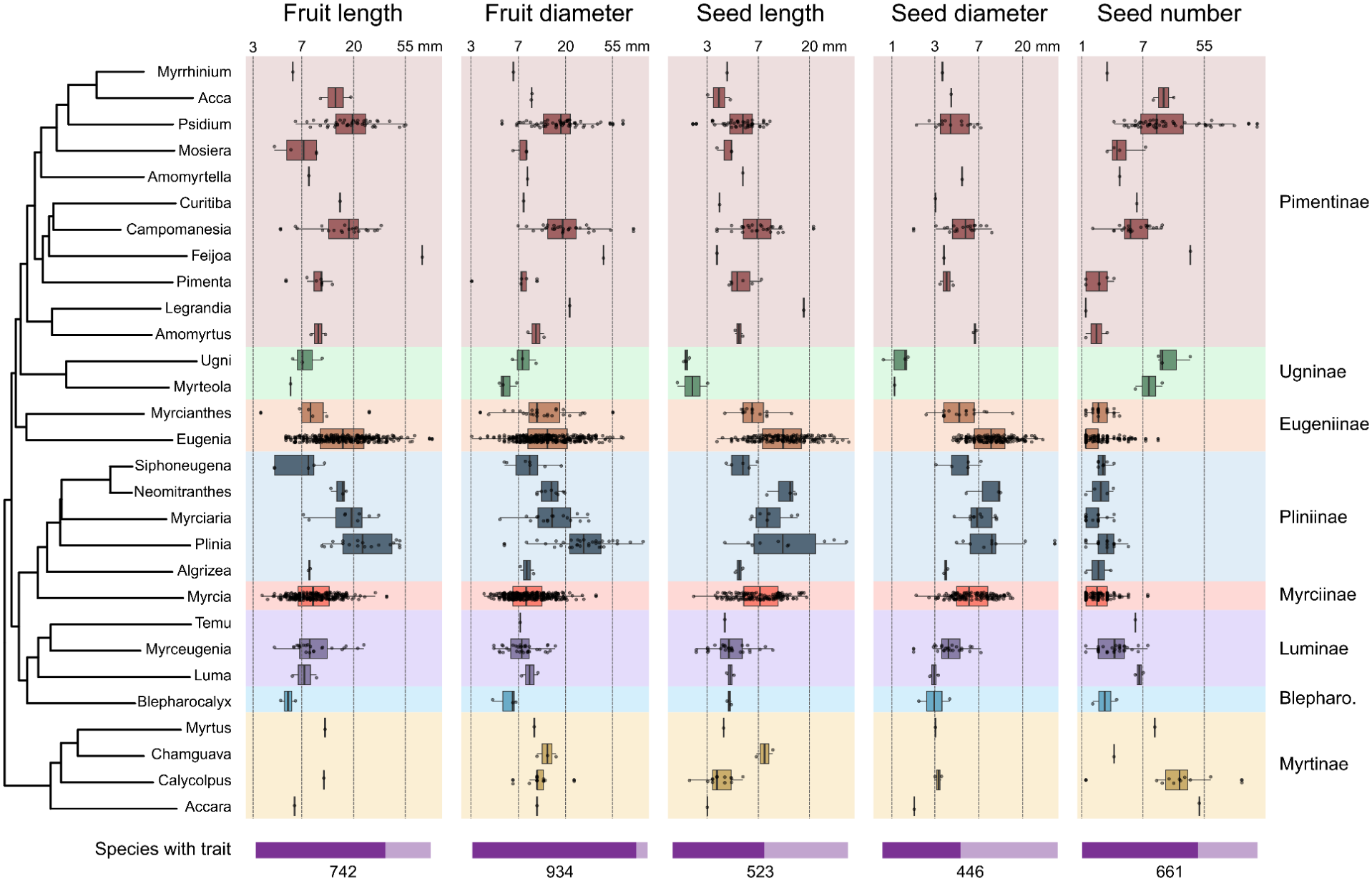
Species mean trait values across Neotropical Myrtaceae genera. For visualization purposes, trait values were plotted on a logarithmic scale, with x-axis expressed in millimeters to facilitate interpretation. Colors represent Myrteae subtribes (Blepharo. = Blepharocalycinae). Purple bars indicate the total number of species in the database with data available for each trait (total database coverage = 997 species and 13 varieties). Genus-level phylogeny is adapted from NMWG et al. (2024)

**Figure 3.**
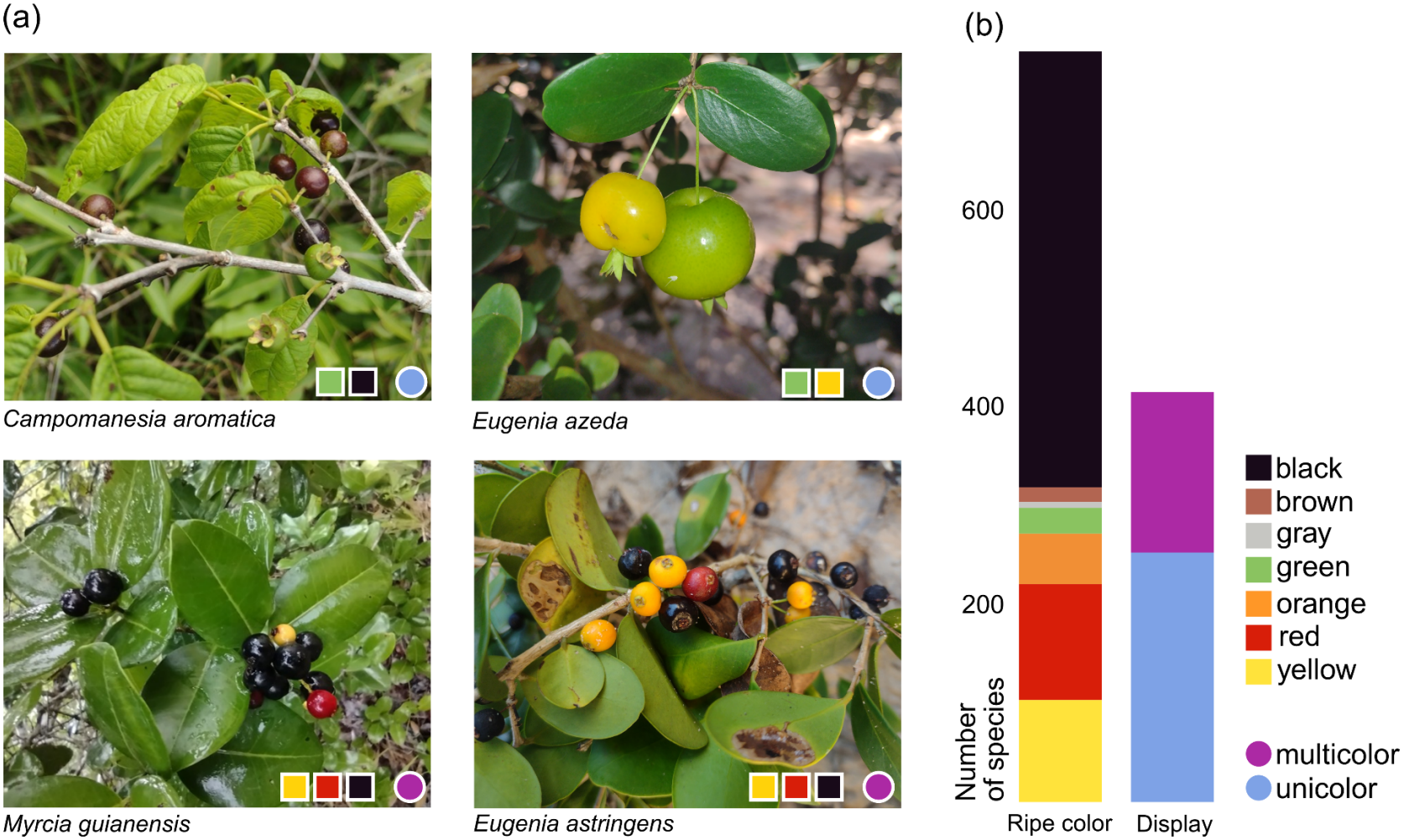
(a) Examples of fruit images of species included in the database (photos: HFC); colored squares indicate fruit color and display type. (b) Number of taxa in the database with information on ripe fruit color and display type.

Data on fruit morphology are available for >90% of the species, in contrast to data on seed morphology that are available for about 50% of the species. The average fruit size is 14.30 ± 9.97 mm in length and 13.29 ± 9.73 mm in diameter, whereas the average seed size is 8.60 ± 5.76 mm in length and 7.37 ± 4.58 mm in diameter. The number of seeds per fruit ranges from 1 to 264, with an average of 5 ± 17 seeds per fruit. Around 30% of the species produce single-seeded fruits. The variation in fruit and seed traits across subtribes and genera is summarized in Fig. 2.

## Summary of methods

Fruit and seed traits were compiled through an extensive Google Scholar search using combinations of terms related to fruits, seeds, morphology, Myrtaceae, and individual species names in English and Portuguese. When publications were unavailable online, authors were contacted directly. The database was continuously updated with literature published from 1857 to August 2024, with most studies published between 2000 and 2024 (*n* = 206, 82.4%). Given the broad temporal coverage, nomenclature was standardized to the taxonomy accepted on July 8, 2026. Brazilian species followed Flora e Funga do Brasil (2026), while non-Brazilian species followed Plants of the World Online (POWO, 2026). Subtribal classification followed Lucas et al. (2019), and infrageneric classifications followed Mazine et al. (2016) for *Eugenia*, Lucas et al. (2018) for *Myrcia*, Proença et al. (2022) for *Psidium*, and Stadnik (2020) for Pliniinae. Species identified as varieties in the original sources were retained as such unless POWO (2026) or Flora e Funga do Brasil (2026) did not accept the variety.

To complement the database, field expeditions were conducted across various regions of Brazil, including the states of Bahia, Distrito Federal, Pará, Paraná, Rio de Janeiro, Rio Grande do Norte, and São Paulo. Between two and ten ripe fruits and their seeds were measured per species. Measurements were also obtained from one to three herbarium specimens per species, with at least two fruits and their seeds measured per specimen using a digital caliper with 0.01 mm precision. For each species, measurements from the available field-collected individuals and herbarium specimens were used to calculate separate species-level means. Each mean was recorded as a distinct database entry, classified as either “Field” or “Herbarium,” analogous to records derived from the literature. To ensure data reliability, Myrtaceae experts (co-authors) reviewed the database, validating the compiled records and supplementing them with field observations for several species—particularly regarding fruit color and display

Color perception and description vary among individuals due to biological and cultural factors (Brown & Lindsey 2004, Jameson 2005, Jaint et al. 2010, Langina-Jansone et al. 2020). Consequently, mature fruits of the same species were sometimes described differently across sources or assigned compound color terms (e.g., yellow-green, reddish-black), complicating single-color standardization. We therefore adopted the following protocol: (1) when only one record was available, the warmest or deepest color was selected (e.g., yellow-green = yellow; reddish-black = black); and (2) when multiple records were available, the majority rule was applied, with the most frequently reported color adopted as the species’ standard. Color records were carefully verified by the authors and corrected when necessary using expert-identified herbarium specimens, field observations, expert knowledge or species descriptions, and consultations with additional taxonomic experts (see Acknowledgements). Literature-based records were checked against field observations, expert knowledge or species descriptions for 611 species (60.5%), iNaturalist photographic records for 14 species (1.4%), and herbarium labels for 51 species (5.0%).

Given the variation in color terminology, similar descriptions were grouped into standardized categories before analysis (Table 1): black, brown, gray, green, orange, red, and yellow. The standardized and corrected color assigned to each species is indicated in the color_pat column. Thus, the final color category may differ from the original descriptions in individual records, reflecting the standardized classification applied across all data sources. To determine the species-level color assignment, we also considered the method of data collection (col_meas_pat). When records from multiple collection methods were available, field observations were prioritized over other sources. When records were available from only one collection method, the final color was assigned based on the most frequent color category among the available records.

**Table 1.** Color standardization of Myrteae fruits.

| **Standard colors** | **Colors described in the original studies** |
|---|---|
| **Black** | alaranjado-atropurpúreo, alaranjado-nigrescente, alaranjado-roxo, alvo-purpúreo, amarelo-vermelho-roxo, amarelo-alaranjado-violáceo-negro, arroxeado, atropurpúreo, atropurpúreo-negro, avermelhado-arroxeado, avermelhado-nigrescente, avermelhado-roxo, azulado-atropurpúreo, azul-escuro-negro, black, blackish-purple-blue, bright red-dark purple, brown-vinaceous, dark, dark purple, dark purple-black, dark red, dark red-black, dark vinaceous-black, dark violet, dark wine-red, deep purple-black, dull purplish, enegrecido, negro, negro-azulado, negro-roxo, negro-roxo-vermelho, negro-vermelho, negro-vináceo, negro-violáceo orange to purple-black, orange-colored-black, pink-red-purple-dark, preto, preto-roxeado, purple, purple to dark-purple, purple-blackish, purplish-black, purplish-dark, purpúreo, púrpuro-enegrecido, púrpuro-negro, púrpuro-violáceo, red or dark, red or purple-blue-black, red to black, red to blackish, red to dark, red-purple, red-black, red-dark reddish, reddish purple, reddish to dark, reddish-dark red, reddish-purple, reddish-vinaceous, red-purple, rosa rosáceo-roxo, rosado, rosado ou roxo–rosado, róseo-roxo, roxo, roxo enegrecido, roxo escuro, roxo ou esbranquiçado, roxo-negro, roxo–preto, roxo-purpúreo, vermelho ou negro, vináceo-roxo, violáceo, violáceo–negro |
| **Brown** | castanho, greenish-brown-maroon, marrom, red-brown, yellowish-brown |
| **Green** | amarelo-esverdeado, esverdeado, verde, verde-amarelado |
| **Grey** | grey, dark grey, light grey, grey to grey–brown, black grey |
| **Orange** | alaranjado, amarelo-alaranjado, laranja, orange, orange-brown, orange-yellow, yellow-orange |
| **Red** | alaranjado ou vermelho, alaranjado-avermelhado, alaranjado-vermelho, amarelado-avermelhado, avermelhado, avermelhado a ferrugíneo-tomentosas, branco-róseo-vermelho, bright red, green–reddish laranja e vermelho-claro, orange to red, orange–red, pink or red, red, red and yellowish, red-yellow, reddish, reddish–white, rosa, vermelho, rosa–choque, rubrae, vemelho, vermelho ou alaranjado, vermelho ou laranja-avermelhado, vermelho–alaranjado, vermelho–amarelado, vermelho–amarelo, vermelho–escarlate, yellow-red, vinaceous, vinho–escuro, vinosos–nigrescentes, vermelho pupúreo, vermelho–arroxeado, vermelho-escuro, vermelho–escuro–negro, vermelho-negro, vermelho–preto, vermelho-roxo, vermelho–vináceo, vináceo, vináceo–negro, vináceo–nigrescente |
| **Yellow** | amarelado, amarelo, amarelo-alaranjado, amarelo-claro, amarelo-esverdeado, amarelo-laranja, bege-creme, dull yellow, greenish-orange-yellowish, green-yellow, ocráceo, ocráceo-pubescente, pale yellow, pale yellow-brownish, verde-amarelado, verde-amarelo, yellow, yellowish, yellowish-green yellowish-orange, yellow-orange |

In Table 1, a single original color descriptor may occasionally correspond to two distinct standardized categories. For example, *Psidium guineense* and *Psidium appendiculatum* were originally described in the literature as “amarelo-esverdeado” (greenish-yellow); however, based on field observations, these records were corrected and standardized as yellow for *P. guineense* and green for *P. appendiculatum*.

Adjustments were made to account for fruit color variation across geographic regions and developmental stages. For instance, *Myrciaria tenella* may produce yellow or red fruits, whereas *Eugenia uniflora* and *Myrciaria floribunda* are typically described as having mature red fruits but may produce black fruits in certain localities. In such cases, the most frequently reported color across all records was adopted. Some *Campomanesia* species (*C. guazumifolia, C. neriiflora*, and *C. schlechtendaliana*) have green fruits at ripening but turn yellow after falling from the tree; therefore, yellow was adopted as the final color. Similarly, when yellow fruits transition to orange after maturation, orange was selected. The gray category was used for fruits whose epicarp is generally black but appears gray because it is densely covered by wax deposits or trichomes, a condition observed in some Myrtaceae species (e.g. *Eugenia mosenii*, and *Myrcia pubipetala*).

Fruit displays were classified following Cunha et al. (2026). Unicolored displays contained two colors simultaneously (i.e. the immature and final ripe fruit colors) whereas multicolored displays contained three or more colors, including intermediate stages during fruit maturation (Fig. 3). Display information was obtained from field observations, literature, photographic records, and herbarium labels. Literature-based records were checked against field observations, expert knowledge, or species descriptions for 338 species (33.5%), iNaturalist photographic records for 25 (2.5%), and herbarium labels for 26 (2.5%). We also recorded the number and identity of colors in each display.

Vegetation types were maintained according to the original research. Biomes follow the Brazilian biome classification and were maintained as in the original research (WWF, 2026). For province classification, we used the system proposed by Morrone et al. (2022). Through this process, we obtained the updated provinces of occurrence for the species in the Neotropics.

## Limitations and Challenges

The database was thoroughly checked and verified. Descriptive statistics and data filtering were used to identify anomalous values, including measurements recorded in cm instead of mm, or vice versa. Whenever possible, these errors were verified and corrected by consulting the original sources and contacting the authors of the relevant references. Unverifiable problematic records were removed. Morphological data from Bufalo et al. (2016) and Bello et al. (2017) were also checked against their original sources and corrected or excluded when discrepancies were found. Consequently, original sources were used whenever available, and only 31 and 57 species records with morphological data were retained from these two reviews, respectively. Data based on immature fruits were excluded. If users identify errors, please notify the authors.

When using these data, users should consider the source of morphological measurements, particularly for fruit length and diameter. Neotropical Myrtaceae generally bear fleshy fruits, whose dimensions may be distorted by pressing and drying during herbarium preparation (Kilsztajn et al., 2026a). Herbarium measurements may therefore not accurately represent fresh fruit dimensions. One way to account for this issue is to use paired fresh and herbarium measurements to build predictive models of fresh fruit dimensions; Kilsztajn et al. (2026a) present such a framework for Neotropical Myrtaceae, with promising results.

Fruit morphology data were more common than seed morphology data. Therefore, we emphasize the need for researchers to include seed morphology in species descriptions whenever possible and during the sampling of fruits. We recommend recording mature fruit color during field collections, as well as noting the presence of temporal displays and their colors. Additionally, collectors should explicitly state on labels whether fruits are mature or immature. Using ‘green fruit’ to mean ‘immature’ is a source of confusion, as some fruits remain green even when fully mature.

## Scientific Opportunities

This dataset provides opportunities to investigate macroecological and macroevolutionary patterns of fruit and seed trait evolution across Neotropical ecosystems. Its broad spatial, temporal, and taxonomic coverage supports tests of the dispersal syndrome hypothesis, trait matching and the relationships between reproductive traits, lineage diversification, historical biogeography, and habitat shifts (e.g. Messeder et al. 2024, Onstein et al. 2017, Cabral et al. 2025, Cunha 2024, Lu et al. 2019, Kilsztajn et al. 2026b, 2026c). The data can also support studies of correlated trait evolution, ancestral state reconstruction, and taxonomic and systematic questions (e.g. Sinnott-Armstrong et al., 2020; Valdemarin, 2023). More broadly, they provide a baseline for assessing the consequences of defaunation (Galetti et al., 2013; Dirzo et al., 2014), mutualistic network disruption, climate change (Fricke et al., 2022), and habitat fragmentation (Fontúrbel et al., 2015), including their effects on functional diversity and conservation priorities (Bello et al., 2015).

## Acknowledgments

We would like to thank all our taxonomist colleagues who helped us by sharing information on taxon nomenclature and identification, species distribution and fruit color (Amélia Tuler, Augusto Giaretta, Bruno Amorim, Duane Lima, Jaquelini Luber, Leidiana Santos, Matheus Santos, Priscila Rosa and Thais Vasconcelos) and all the authors of the publications used here who were willing to revise the original data when doubts arose. HFC received two years (2019-2021) of funding of scientific initiation scholarships from PIBIC-UFRN (Programa Institucional de Bolsas de Iniciação Científica). YK thanks the Sociedade Botânica do Brasil (SBB) for financial support through the Scientia Amabilis 2024 grant. CEBP thanks Conselho Nacional de Desenvolvimento Científico e Tecnológico - Brazil (CNPq) for her PQ2 *Produtividade em Pesquisa* Grant *Taxonomia, Ecologia e Conservação das Myrtaceae do Cerrado e Floresta Atlântica do Nordeste*. KSV thanks Bolsas FUNBIO – Conservando o Futuro, and the São Paulo Research Foundation (FAPESP), Brazil (#2022/12597-0 and #2025/21381-0). This study was financed in part by the Coordenação de Aperfeiçoamento de Pessoal de Nível Superior - Brasil (CAPES) - Finance Code 001 and by the Instituto Serrapilheira (grant number Serra-R-2111-39858).

## Notes

**Conflict of Interest statement** None declared

### Competing Interest Statement

The authors have declared no competing interest.

